# Adaptive meiotic drive in selfing populations with heterozygote advantage

**DOI:** 10.1101/666180

**Authors:** Evgeny Brud

## Abstract

The egalitarian allotment of gametes to each allele at a locus (Mendel’s law of segregation) is a near-universal phenomenon characterizing inheritance in sexual populations. As exceptions to Mendel’s law are known to occur, one can investigate why non-Mendelian segregation is not more common using modifier theory. Earlier work assuming sex-independent modifier effects in a random mating population with heterozygote advantage concluded that equal segregation is stable over long-term evolution. Subsequent investigation, however, demonstrated that the stability of the Mendelian scheme disappears when sex-specific modifier effects are allowed. Here I derive invasion conditions favoring the repeal of Mendelian law in mixed and obligate selfing populations. Oppositely-directed segregation distortion in the production of male and female gametes is selected for in the presence of overdominant fitness. The conditions are less restrictive than under panmixia in that strong selection can occur even without differential viability of reciprocal heterozygotes (i.e. in the absence of parent-of-origin effects at the overdominant fitness locus). Generalized equilibria are derived for full selfing.

## Introduction

Despite a growing number of examples where the rule of Mendelian segregation is violated (e.g. Fishman and Saunders 2008, Didion et al. 2016, Nuckolls et al. 2017), exceptions to fair meiosis in the form of meiotic drivers are thought to be relatively uncommon (Crow 1991). Among the exceptions, many cases of meiotic drive may be transient in that they involve the spread of an initially rare segregation distorter that quickly rises all the way to fixation. Drivers of this sort are consistent with the observation of widespread adherence to the Mendelian scheme, since fixation entails the loss of manifestation of the drive phenotype. Other cases of meiotic drive may involve homozygous fitness costs of sufficient magnitude to allow for the initial increase of a driver but not its fixation, and so give rise to a condition of persistent variation for segregation distortion as is found in mice (e.g. t-haplotypes; Dunn 1957), Drosophila (e.g. SD-haplotypes; Sandler, Hiraizumi, and Sandler 1959), the chromosomal knobs of maize (Rhoades 1942), and various cases of sex chromosome drive (e.g. Gershenson 1928), among a number of other examples; often such drivers carry a number of fitness costs in both heterozygous and homozygous genotypes (reviewed in Burt and Trivers 2006). But even these cases are unstable in the long run, since an intermediate frequency of a selfish driver will select for suppressors throughout the genome, and so the existence of segregation distortion at some locus is regularly destabilized by adaptive countermeasures aimed at restoring the Mendelian order.

Considerations such as these seem to argue that Mendelian ratios ought to be widespread (in accord with observation) and evolutionarily stable. However, Úbeda and Haig (2005) proved the existence of conditions in which rare modifiers are expected to invade a resident population by virtue of promoting meiotic drive at unlinked autosomal loci subject to fitness variation. In the present work, I demonstrate that the conditions for evolving a stable phenotype of meiotic drive are even less restrictive when self-fertilization is included in the model.

The general evolutionary causes of autosomal adherence to Mendelian segregation has received sporadic attention in the theoretical population genetics literature since it was first modeled in the 1970s (Hartl 1975; Liberman 1976, 1990; Thomson and Feldman 1976; Liberman and Feldman 1980, 1982; Lloyd 1984; Eshel 1985; Úbeda and Haig 2005, Brandvain and Coop 2015). One branch of these efforts assumes a random mating population with a focal locus subject to di-allelic variation and heterozygote advantage, in which the alleles have evolved to a stable equilibrium. A second locus is assumed to be variable for alleles that modify the segregation ratio at the focal fitness locus. The key question is: what are the conditions favoring the spread of a rare modifier? Early models investigated sex-independent variables and parameters. Notable results include Liberman (1976), who found that in a population fixed for a resident phenotype of Mendelian segregation, rare modifiers could spread under broad conditions: any rate of recombination less than one-half can result in the invasion of a drive-enhancer (i.e. a modifier that imposes or increases a deviation from equal segregation) given that it is sufficiently strong, with tight linkage being especially conducive to invasions. Eshel (1985) showed that in the case of free recombination, the only kinds of modifiers which can invade are those that reduce the intensity of drive; enhancers are uniformly selected against.

These results recall the metaphor of Leigh (1977) who likened the genome to “a parliament of genes” which enforces behavior consistent with the common good against “‘cabals of a few’ conspiring for their ‘selfish profit’” (p. 4543). Modifiers with sufficient linkage to a distorting locus can form a selfish cabal, but the presumably greater number of unlinked modifiers act to police outlaw behavior.

An unsatisfying aspect of the work mentioned above is the assumption of sex-independent effects. If a sex-differentiated population is assumed, and if accordingly modifiers have sex-specific effects on male and female segregation, then an analysis of modifier invasion demonstrates that unlinked genes should evolve to subvert Mendelian ratios after all (Úbeda and Haig 2005). The benefit in doing so owes to the assumption that a modifier acts on a locus exhibiting heterozygote advantage. Oppositely-directed segregation schemes, in which an allele has a segregation advantage in one sex but a segregation disadvantage in the other sex, are selected for as a mechanism of producing a super-Mendelian proportion of fit heterozygous offspring, and thus evolve to reduce the genetic load caused by the production of unfit homozygous progeny (segregation load). This possibility was absent in the sex-independent models. On the assumption of symmetric overdominance, in which parent-of-origin effects on fitness are absent (i.e. “Aa” and “aA” genotypes have identical fitnesses, where the order of the alleles distinguishes parental origin), rare modifiers of the segregation scheme are selected to increase at a slow arithmetic rate. Asymmetric overdominance in fitness, in which “Aa” and “aA” genotypes differ but their average fitness is greater than either homozygote, is a situation which selects for the repeal of Mendelian law at a speedy geometric rate. Úbeda and Haig (2005) discovered this phenomenon, which I refer to as “adaptive meiotic drive” in order to contrast such cases with selfish distortions of meiosis; the concept is defined here as any distortion of segregation ratios that reduces the segregation load in a population, relative to the Mendelian expectation. Importantly, the phenotypic state characterized by long-term stability is the rather un-Mendelian scheme of “all-and-none segregation”, in which ratios are maximally distorted in male and female meiosis, but in opposite directions (see also Úbeda and Haig 2004).

The importance of the sex-specific results with respect to biological phenomena is unclear given that asymmetric overdominance in fitness is likely a rare class of balanced polymorphism. Such a fitness scheme involves the simultaneous realization of heterozygote advantage and parent-of-origin effects (e.g. genomic imprinting), examples of which are lacking. Another factor constraining the spread of adaptive drive is that with respect to symmetric overdominance, the repeal of the Mendelian process is associated with a slow rate of modifier invasion that can easily be counteracted by associated costs to reproduction (e.g. male fertility reduction). Nevertheless, their results reveal that the reasons for the ubiquity of Mendelian segregation are more obscure than is generally appreciated (Úbeda 2006).

The impact of inbreeding on the invasion of unlinked modifiers of drive in the context of heterosis has not previously been investigated, and it is reasonable to suspect that the mating system will have a qualitative impact on the evolutionary dynamics of adaptive meiotic drive. Here I investigate a model that extends the conditions under which Mendelian segregation is selected against to the case of partially and fully selfing populations. Invasion conditions are derived for unlinked modifiers that impose a mutant segregation scheme in the context of symmetric overdominance in fitness. Likewise, I assess the long-term stability of all-and-none segregation to unlinked modifiers that alter the ratios closer to equality. In addition, I derive generalized equilibrium genotype frequencies for the case of full selfing.

## Model

Assuming a monoecious population with sex-differentiated meiosis, I examine a two-locus, two-allele discrete-generation deterministic model with an autosomal fitness locus, *A*, and an unlinked autosomal modifier locus, *B*. The fitness scheme is W_Aa_ = 1 > W_AA_, W_aa_ ≥ 0. In heterozygotes, the resident segregation ratio of *A*: *a* is equal to 1-*k*_*m*0_: *k*_*m*0_ in sperm and 1-*k*_*f*0_· *k*_*f*0_ in eggs of wild-type BB individuals, and in the mutant genotypes Bb or bb, the ratio is 1-*k_m_*: *k_m_* in sperm and 1-*k_f_*: *k_f_* in eggs. Part of the population (1-*S*) are the offspring resulting from outcrossing, assuming random union of gametes. The remainder (*S*) owe to diploid selfing. With 10 possible two-locus diploid genotypes, the dynamical equations for the genotypic proportions are given by:

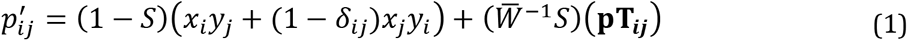

which is valid for *i, j* = 1, 2, 3, 4 and such that *ij* and *ji* are not distinguishable. 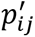 is the next-generation frequency of the genotype composed of haplotypes *i* and *j* (the labels are AB = 1, Ab = 2, aB = 3, ab = 4). The *x_i_* and *y_j_*terms are the male and female gametic frequencies, respectively, subsequent to zygotic selection of diploids, and are given by 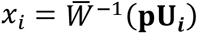 and 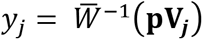, where 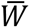 is the mean population fitness, equal to Σ Σ *P_ij_*W_*ij*_. **p** is a row vector of genotype frequencies prior to zygotic selection, and such that Σ Σ *P_ij_* =1. W_*ij*_ is the zygotic fitness of genotype *ij* and is equal to W_AA_ for *ij* = 11, 12, 22 and is equal to W_aa_ for *ij* = 33, 34, 44; all other subscripts correspond to heterozygotes associated with a fitness of 1. **U_*i*_** and **V_*j*_** are the *i*^th^ and *j*^th^ column vectors from the 10×4 matrices (**U** and **V**) governing transmission and selection in outcrossers. *δ_ij_* is the Kronecker delta. **T_*ij*_** is a column vector corresponding to the production of genotype *ij* from the 10×10 matrix (**T**) describing transmission and selection in selfers. **U, V**, and **T** are presented in the appendix.

## Results

### External stability of a Mendelian partial selfing population to adaptive meiotic drive

Assuming a population is initially fixed for the Mendelian-favoring BB genotypes, variation at *A* will be determined by the forces of selection against homozygotes and the rate of selfing. Equilibrium properties for single-locus systems of selfing and overdominant fitness have been investigated previously (Hayman 1953, Workman and Jain 1966). The state space for a single-locus partial selfing population with two alleles is fully described by the allele frequency (q) and the inbreeding coefficient (F). One can formulate a model in which *x* = *p*^2^ + *Fpq; y* = 2*pq*(1 – *F*); *z* = *q*^2^ + *Fpq*, where the frequency of *A* is *p* = 1 – *q*, and the frequencies of AA, Aa, and aa are *x,y*, and z, respectively. (Aa and aA are not distinguished). The influences of selection, selfing, and Mendelian segregation are such that in the next generation genotypic frequencies are

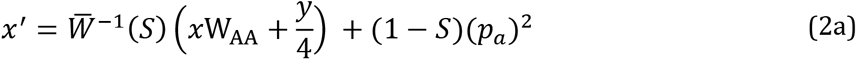

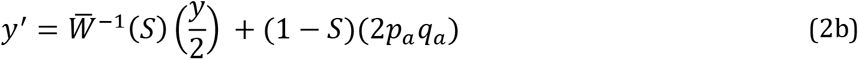

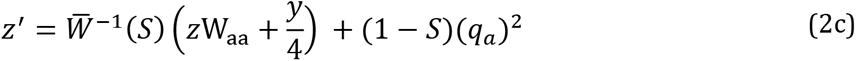

where 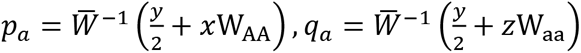 and 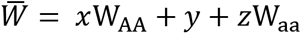 are respectively the frequency of *A* and *a* alleles among outcrossers after selection (*p_a_, q_a_*) and 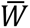 is the mean fitness of the population. Since 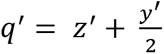 and 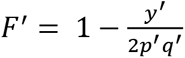 are the frequency of the *a* allele and the inbreeding coefficient in the next generation, setting the change in *q* and *F* over successive generations equal to zero yields a pair of equilibria in 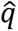 and 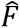, only one of which is biologically feasible in the interior of allele frequency space (i.e. such that both 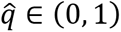 and 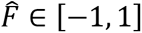 hold true). For tractability of the stability analyses, I assume equal fitness of the homozygotes, *W_AA_* = W_aa_ = W_*G*_, which simplifies the equilibrium so that 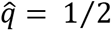 and 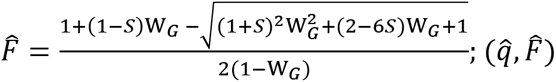 is internally stable 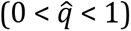 for any rate of partial selfing and any intensity of selection against homozygotes (Supplemental file). Stability under full selfing requires a restrictive condition, namely a greater than two-fold heterozygote advantage (Supplemental file). Treatment of modifier invasion under full selfing is postponed for a later section (**Result 3**).

The stability of 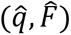 to the introduction of infinitesimal variation at *B* (external stability) can be ascertained by evaluating the Jacobian at equilibrium, which yields an upper triangular block matrix, 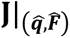. The submatrices on the diagonal are **J_res_**, a 3×3 matrix corresponding to a resident population variable only at *A*, and **J_mut_**, a 7×7 matrix corresponding to the dynamics of a rare mutant at *B* (Supplemental file). The eigenvalues of **J_res_** are less than one in magnitude, and so stability is determined by the leading eigenvalue (*λ_L_*) of **J_mut_**; if |*λ_L_*| > 1, then a rare *b* allele invades at a geometric rate (Otto and Day 2007). Assuming a resident population characterized by Mendelian ratios (i.e. 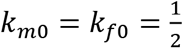):

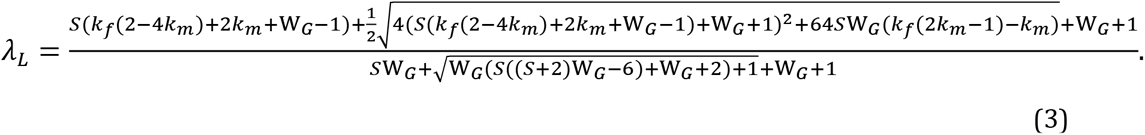

**Result 1**: The parameters for which |*λ_L_*| > 1 holds true are broad. Resident Mendelian populations with partial selfing (0 < S < 1) and heterozygote advantage (0 ≤ W_*G*_ < 1) select for modifiers that impose oppositely-directed sex-specific meiotic drive of any intensity:

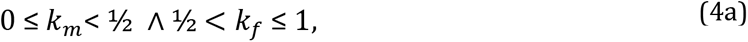

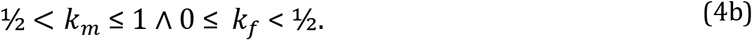

While the invasion analysis provides the conditions for the long-term spread of a rare modifier, it is of interest whether the time course of modifier evolution is appreciable with respect to the lifespan of a balanced polymorphism. Overdominance in the deterministic model results in a permanent stable equilibrium, but in nature, balanced polymorphisms only last as long they are not replaced by fitter genotypes in the course of environmental changes or evolution of the genetic background (e.g. the resolution of overdominance by gene duplication; Spofford 1969). Therefore, determining whether modifier evolution occurs on a reasonable timescale is a necessary prerequisite to evaluating the importance of the adaptive meiotic drive phenotype. Below, I numerically iterate cases of evolution over a time span of 10^5^ generations for a sample of parameters.

### Numerical iterations of adaptive drive-enhancer invasion

Numerical iterations of the two-locus model with strong magnitudes of drive and selection (assuming equal fitness of homozygotes and perfectly sex-reflected segregation schemes, *k_m_* = 1-*k_f_*) are illustrated in Fig.1 and demonstrate fairly broad invasibility of drive-enhancing modifiers starting from a frequency of 0.01% rising to “fixation” (> 99%) within 10^5^ generations, which often occurred except for the weakest case of selection and drive depicted (i.e. 60:40 drive with 1% fitness reduction). A low selfing rate (S = 2.5%) is seen to be uniformly associated with the slowest rate of evolution; multiple cases of modifier fixation are observed for low selfing nonetheless. As is intuitive, higher intensities of distortion and stronger selection against homozygotes are each associated with quicker fixation of the drive enhancer.

**Fig 1.**
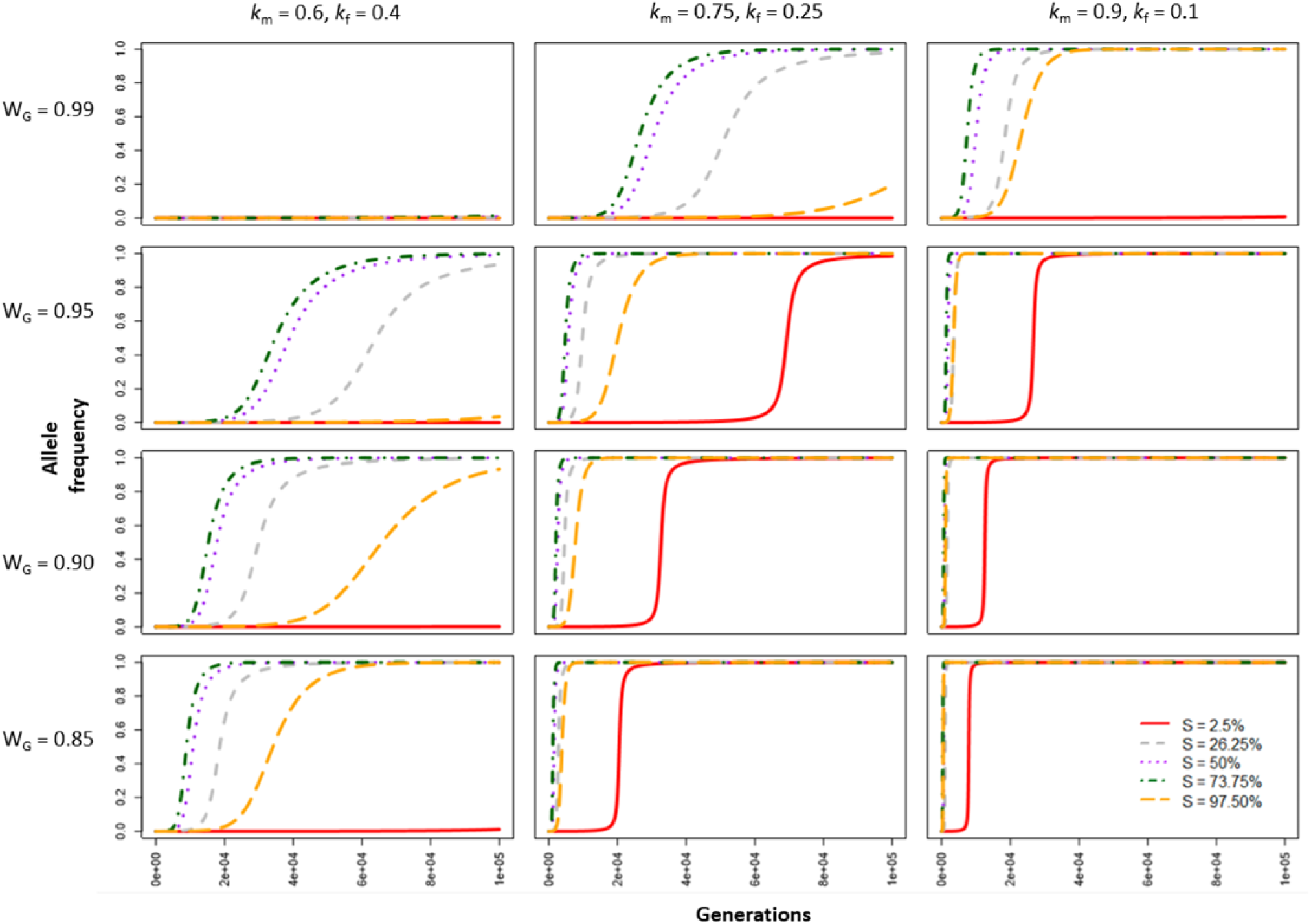
Evolutionary trajectories of the *b* allele for a sample of ‘strong’ parameter combinations. Evolution in these trajectories is depicted over 10^5^ generations, starting from an initial frequency of 10^-4^ for the modifier mutant and linkage equilibrium with the fitness locus *A*. Alleles at the *A* locus remain at a frequency of 50% throughout due to symmetry of the parameter values. For a particular panel, homozygous fitness values are given by the row label and segregation ratios by the column label.

The lack of appreciable modifier evolution within 10^5^ generations for the weakest intensities of drive and selection in Fig. 1 calls for an investigation into cases where only one of these processes exhibits small values, while the other process is strong. An examination of cases reveals that weak segregation distortion and strong selection does not guarantee timely evolution of modifiers (Fig. 2), nor does weak selection and strong distortion (Fig. 3). Nevertheless, with coefficients of sufficient magnitude, cases of drive enhancer fixation are apparent for various levels of selfing. It should be noted that in general, the relation between the selfing rate and the pace of modifier evolution is nonmonotonic; an expression for the selfing parameter that maximizes the rate of modifier invasion can be derived analytically, in terms of W_G_ and *k*, where *k* = *k_m_* = 1-*k_f_* (Supplemental file, Fig. S1).

**Fig 2.**
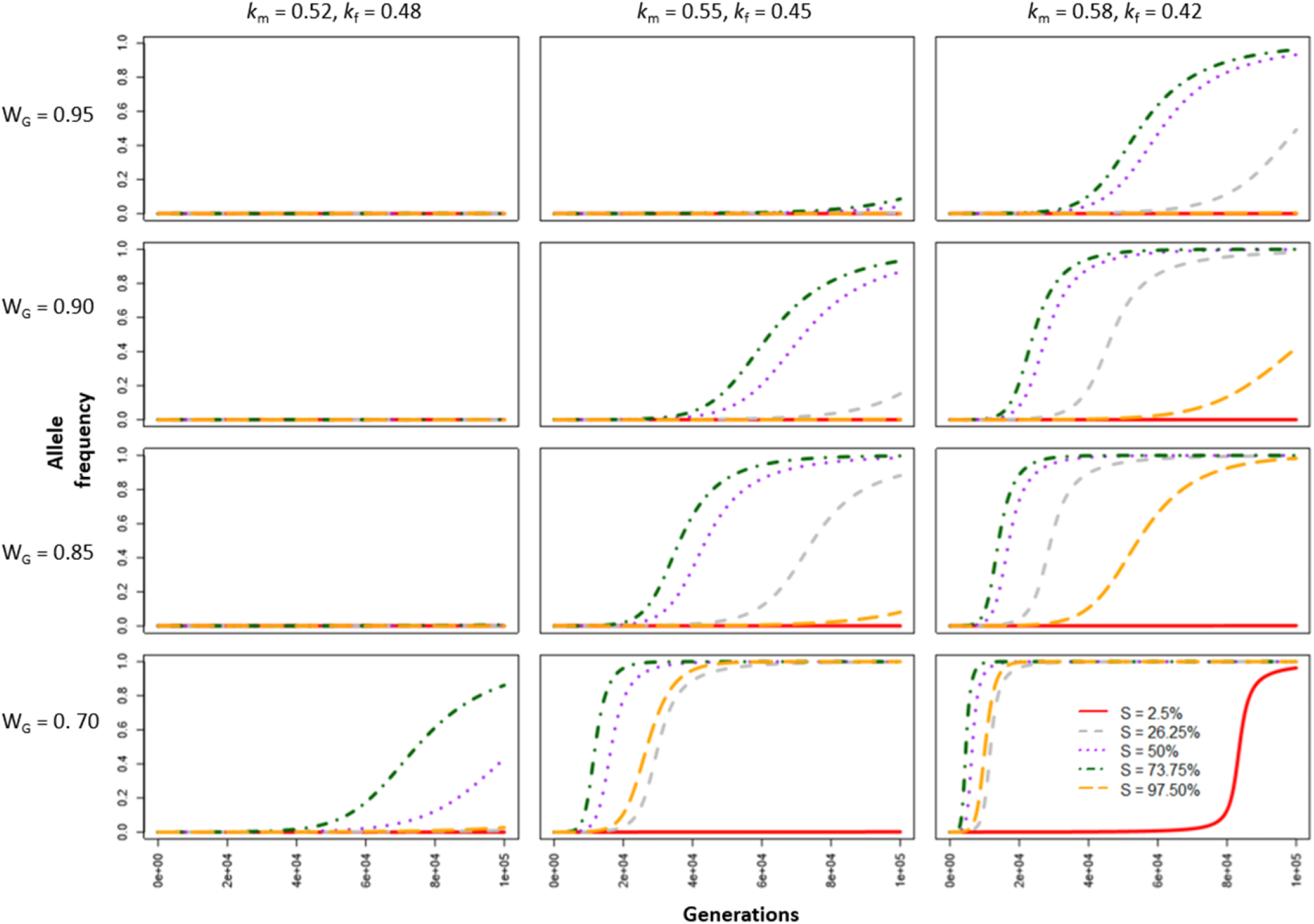
Evolutionary trajectories of the *b* allele for a sample of parameters with strong selection and ‘weak’ segregation distortion. Evolution in these trajectories is depicted over 10^5^ generations, starting from an initial frequency of 10^-4^ for the modifier mutant and linkage equilibrium with the fitness locus *A*. Alleles at the *A* locus remain at a frequency of 50% throughout due to symmetry of the parameter values. For a particular panel, homozygous fitness values are given by the row label and segregation ratios by the column label.

**Fig 3.**
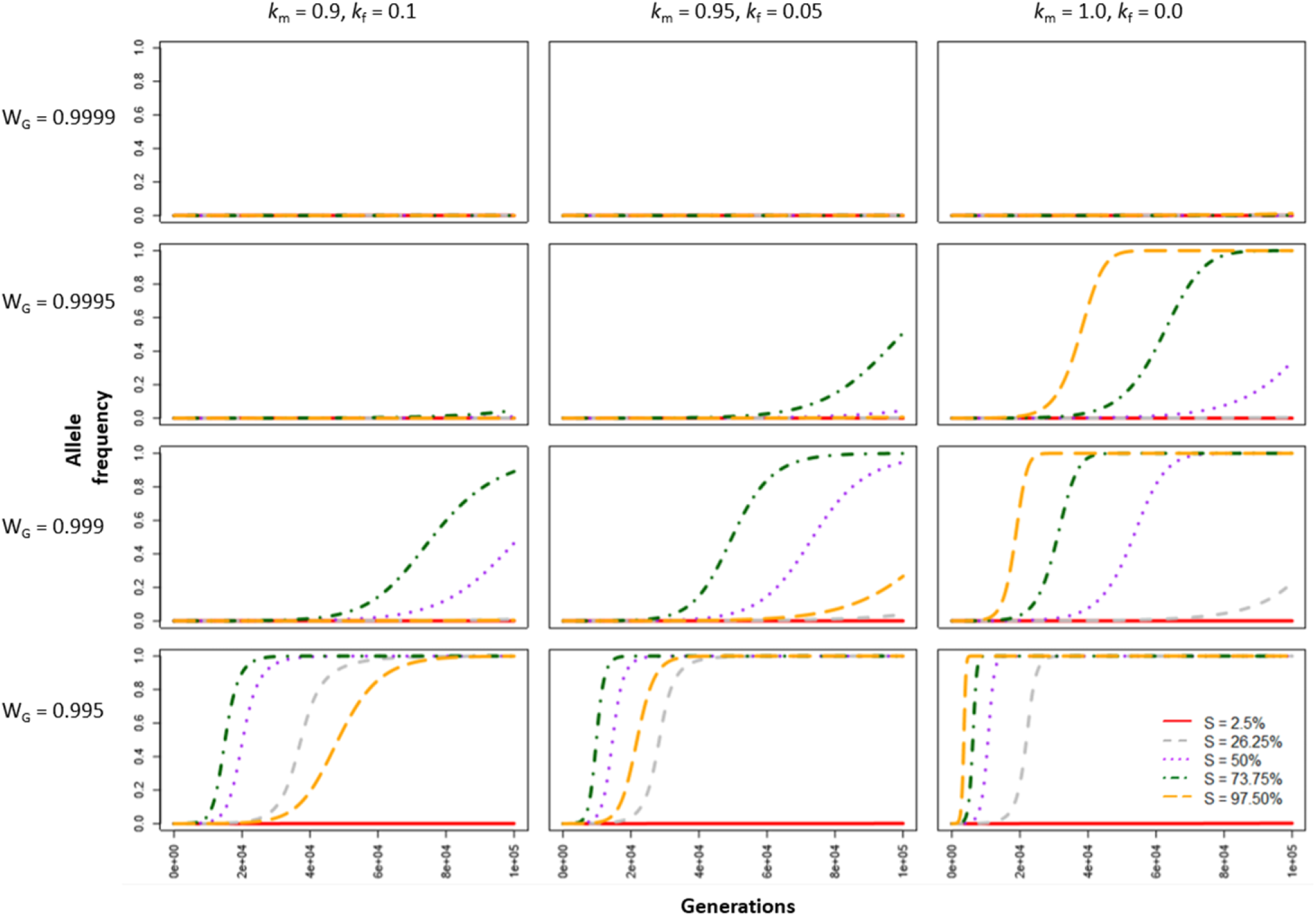
Evolutionary trajectories of the *b* allele for a sample of parameters with ‘weak’ selection and strong segregation distortion. Evolution in these trajectories is depicted over 10^5^ generations, starting from an initial frequency of 10^-4^ for the modifier mutant and linkage equilibrium with the fitness locus *A*. Alleles at the *A* locus remain at a frequency of 50% throughout due to symmetry of the parameter values. For a particular panel, homozygous fitness values are given by the row label and segregation ratios by the column label.

The invasion analysis (**Result 1**) and the preceding numerical results took on the restrictive assumption that homozygote fitnesses were equal, and so I also present a sample of numerical results where this assumption is relaxed. A difference in fitness between the homozygotes (AA and aa) of course results in lower initial heterozygosity at A, and a slower invasion rate of the modifier as compared to the case of equal homozygote fitness (Fig. 4a,b); a shift in the frequency at the *A* locus occurs away from its Mendelian equilibrium to a new value. In addition, if asymmetrically sex-reflected ratios (i.e. *k_m_*≠*k_y_*) are investigated, a modifier polymorphism can appear (Fig. 4c-d). In some cases (Fig. 4c), the mutant modifier winds up at an internal neutral equilibrium because the process of modifier selection causes a frequency shift at the *A* locus into a fixation state; such an event stops any further directional selection at the *B* locus. In other cases (Fig. 4d), alleles at *A* and *B* are both internally stabilized; for the *B* locus, this owes to a balance of forces between (1) an adaptive reduction in the segregation load (i.e. lowered homozygote production) and (2) a maladaptive imposition of a drive load caused by the frequency shift at A (Úbeda and Haig 2005). Yet other cases of asymmetric drive cause a shift at *A* but still allow for the modifier mutant to fix (Fig. 4e,f).

**Fig. 4.**
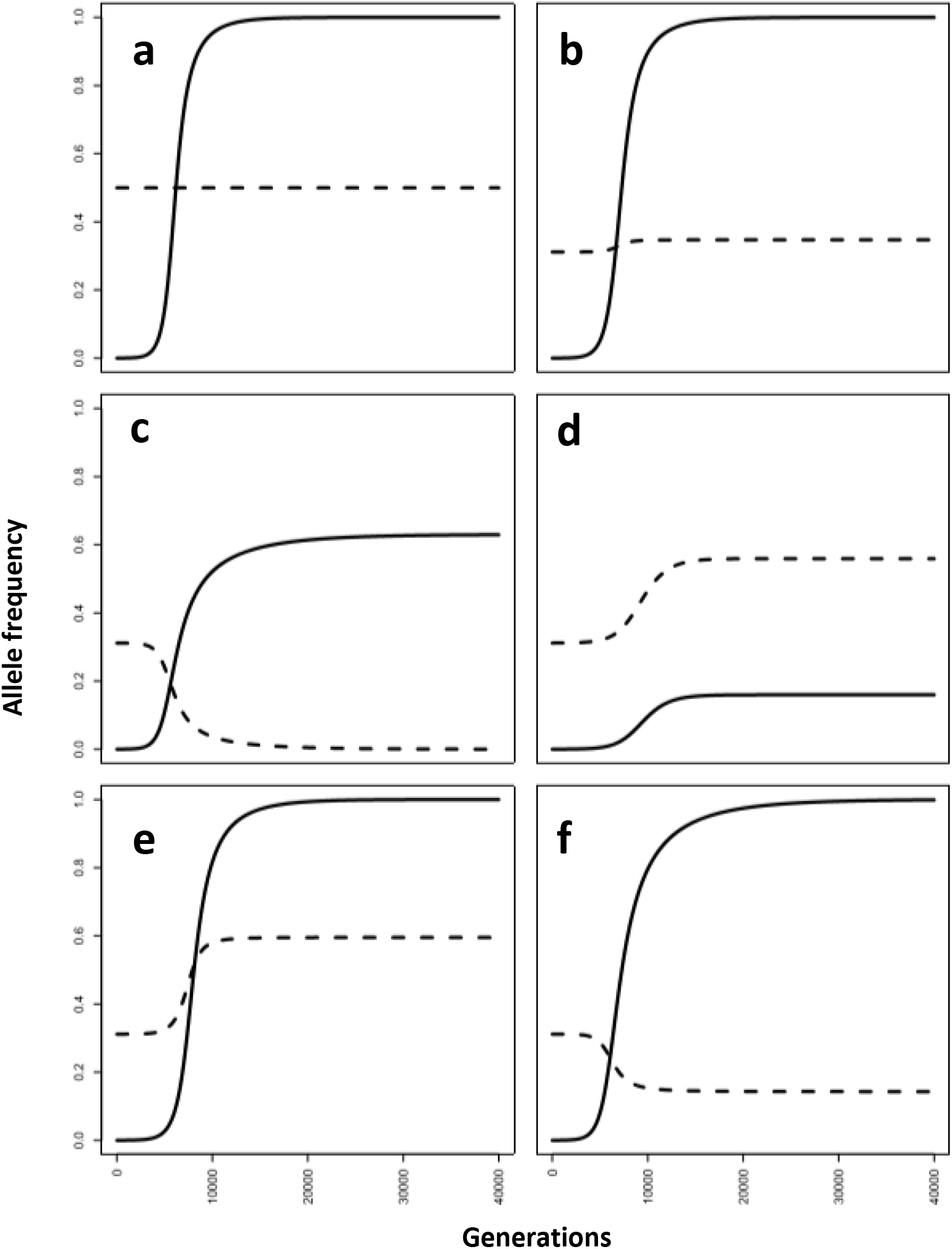
Evolutionary trajectories of adaptive drive evolution with asymmetric homozygous fitnesses and segregation schemes. (Panels a-f) Modifier “b” allele (solid line) is introduced into a Mendelian population at a frequency of 10^-4^ and the “a” allele (dashed) starts at the deterministic equilibrium for single-locus heterosis under equal segregation; change over 40,000 generations is displayed. S = 0.37 and *k_m_* = 0.77 throughout. (a) Equal homozygous fitness and a perfectly sex-reflected segregation scheme (i.e. *k_m_* = 1-*k_f_*): W_AA_ = W_aa_ = 0.95, *k_f_* = 0.23. (Panels b-f) Asymmetric homozygous fitnesses: W_AA_ = 0.962, W_aa_ = 0.938. (b) *k_f_* = 0.23. (c) *k_f_* = 0.19. (d) *k_f_* = 0.315. (e) *k_f_* = 0.255. (f) *k_f_* = 0.21.

### External stability of all-and-none to drive suppressors in a partial selfing population

Úbeda and Haig (2005) reported that all-and-none segregation is stable to mutants of the segregation scheme in random mating populations. I investigate whether all-and-none exhibits stability under partial selfing. The dynamical equations now assume k_m0_ = 1 and k_f0_ = 0 (i.e. one of the two possible all-and-none phenotypes; k_m0_ = 0 and k_f0_ = 1 would do just as well). With equal homozygote fitnesses, the single-locus equilibrium associated with all- and-none segregation, selfing, and overdominance can be shown to equal 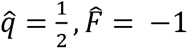, which is an internally stable equilibrium consisting entirely of AaBB genotypes (Supplemental file). The eigenvalues of **J_mut_** in this case are:

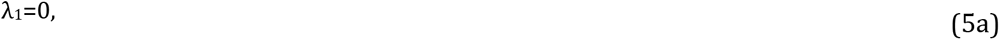

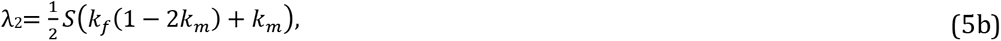

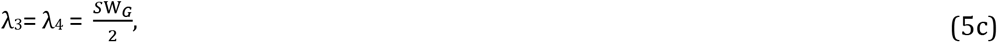

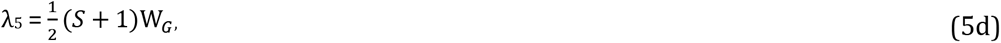

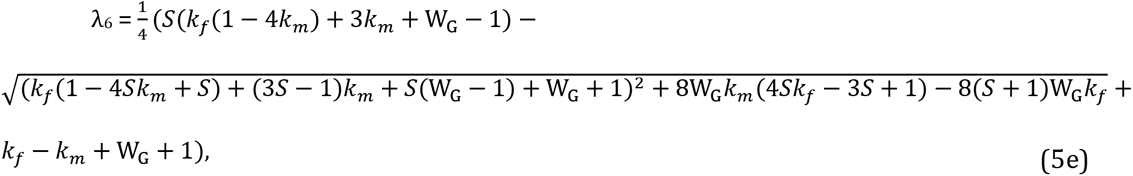

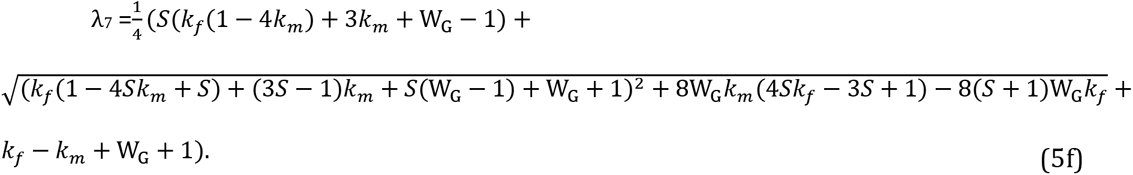

**Result 2**: None of the eigenvalues of **J_mut_** for an all-and-none resident phenotype (Eq. 5a–f) are ever greater than one in magnitude in a partial selfing population. λ_7_ takes the unit value when the mode of all-and-none segregation mutates from (*k*_*m*0_ = 1, *k*_*f*0_ = 0) to (*k_m_* = 0, *k_f_* = 1). Otherwise, all eigenvalues are less than one. This demonstrates the long-term stability of all-and-none segregation to any mutants of the segregation scheme in a partial selfing population with symmetric overdominance and equal fitness of homozygotes.

As an extension of **Result 2**, it can be shown that the direct invasion of Mendelian segregation (*k_m_* = 1/2, *k_f_* = 1/2) into a resident all-and-none population cannot occur for generalized values of W_AA_ and W_aa_ (i.e. with no constraint on their equality); the eigenvalues are complicated (Supplemental file). Similarly, the direct invasion of a sex-limited Mendelian ratio (and maintenance of maximal distortion in the other sex) is also selected against in these more general circumstances (Supplemental file).

### The case of full selfing (S = 1)

In the absence of outcrossing, the model reduces to a linear dynamical system for which the equilibrium can be readily derived for any initial genotype frequencies and general parameters. Following Karlin (1968), given a 10×10 matrix (**T**) that transforms the row vector of two-locus diploid genotype frequencies (**p**) on a generation-by-generation basis, the time course of evolution can be written as: 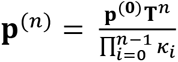, where *n* is the number of generations, *κ* is a normalizing constant that ensures the elements of **p** add up to one in each generation, and **p**^(**0**)^ is the vector of initial values. As *n* → ∞, the vector **p**^(*n*)^ converges to the equilibrium vector **M**. If an adaptive drive-enhancing modifier is introduced into a population at stable equilibrium owing to overdominance (i.e. 0 ≤ W_*AA*_, W_*aa*_ < 1/2), then the leading eigenvalue of **T** is *λ* = *k_f_* + *k_m_* – 2*k_f_k_m_*, which has multiplicity 1. The vector **p**^(*n*)^ as *n* → ∞ can be written as 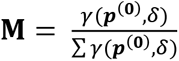 where *γ* and *δ* are respectively the left and right eigenvectors associated with λ, normalized to be biorthogonal, and (**p**^(**0**)^, *δ*) is the inner product.

**Result 3**: Under full selfing, the equilibrium two-locus genotype frequencies are

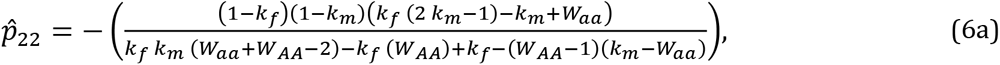

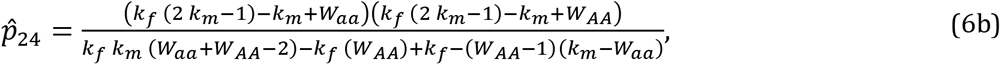

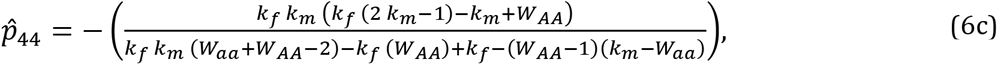

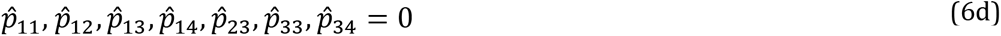

which correspond to positive frequencies for only the Ab/Ab, Ab/ab, and ab/ab genotypes, indicating globally convergent fixation of an adaptive drive-enhancing modifier for the generalized case.

All invading adaptive drive-enhancers converge to fixation, even for asymmetrically-reflected drive schemes, and so there are no conditions for stable or neutral modifier polymorphisms as under partial selfing. With the establishment of sufficient distortion, equilibria are characterized by negative inbreeding coefficients owing to an excess of heterozygotes, relative to panmictic Hardy-Weinberg proportions, even though selfing is maximal (Fig. 5). Upon drive-enhancer fixation, the only modifiers which can subsequently invade are those that impose even stronger oppositely-directed distortion. All-and-none segregation under full selfing and stable heterozygote advantage is therefore expected to persist over long-term evolution.

**Fig. 5.**
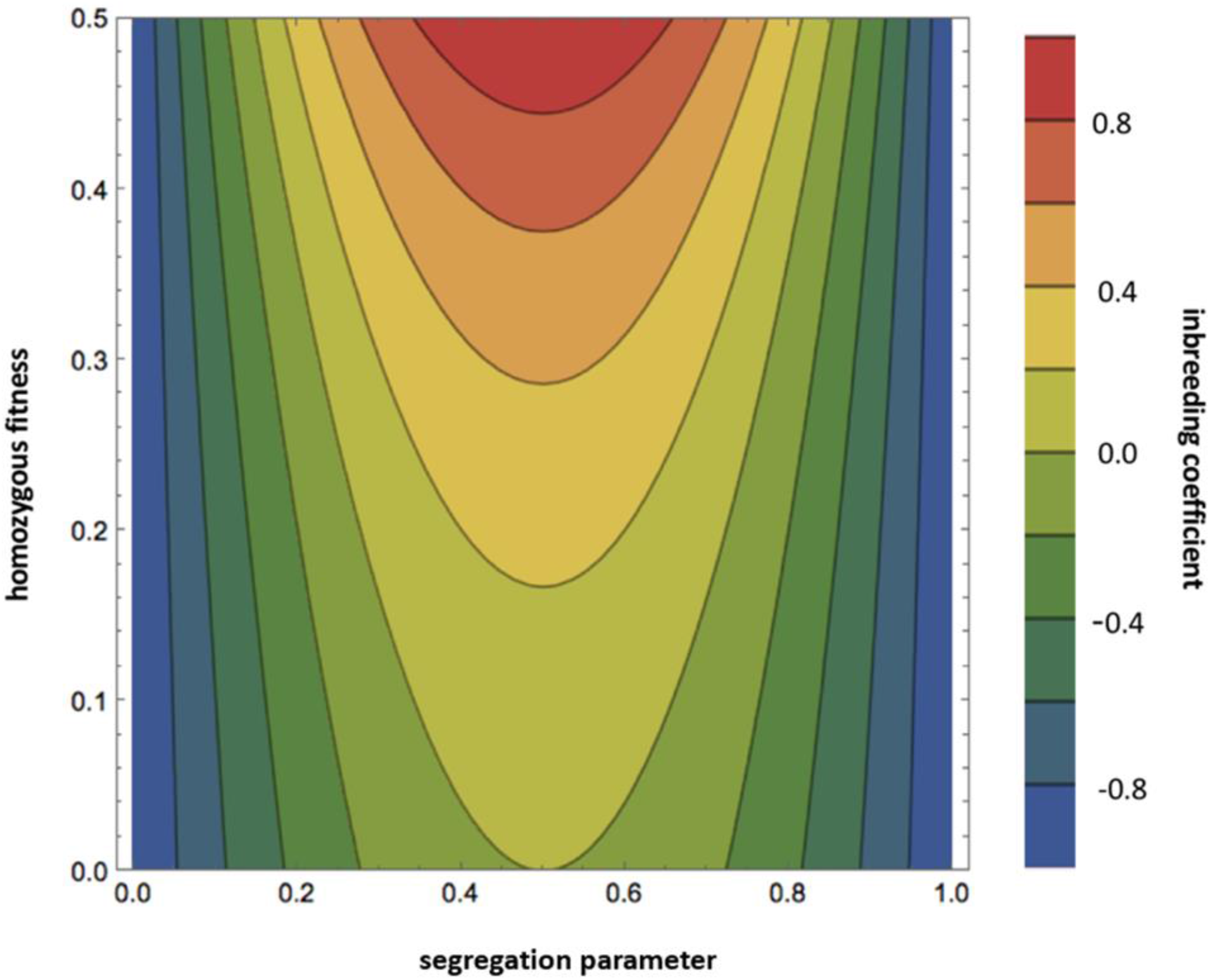
Equilibrium inbreeding coefficients 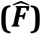 under symmetric homozygote disadvantage, perfectly sex-reflected segregation ratios (*k_m_* = 1-*k_f_*), and obligate selfing. Contours demarcate changes in 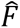 at intervals of 0.2. The parameter space includes only homozygous fitnesses of <50%. Areas of darker green and blue correspond to negative inbreeding coefficients at equilibrium: 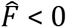 whenever W_G_ < (1-2*k*)^2^. Due to parameter symmetries assumed in this plot, 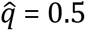 throughout.

## Discussion

The evolutionary instability of the Mendelian scheme on theoretical grounds is seen to owe to the tendency of overdominance to select for sex-specific segregation-distorting phenotypes. While earlier work found that adaptive drive under symmetric overdominance is selected for at a slow arithmetic rate in panmictic populations (the leading eigenvalue corresponding to modifier invasion being at most 1 in magnitude; see Úbeda and Haig 2005, where second-order terms of a Hessian matrix were analyzed in order to ascertain invasion conditions), the inclusion of non-random mating in the model clearly transforms the pace of modifier invasion to that of geometric increase (modifier invasion with a leading eigenvalue greater than 1; **Result 1**, present article). Under panmixia, two mutant individuals would need to mate with each other in order to produce a super-Mendelian proportion of heterozygous progeny, but under mixed or obligate selfing, overproduction of heterozygotes is an immediate consequence of a single mutant individual. In other words, modifier effects on the fitness of selfers are first order since the expression of a fitness advantage does not depend on mating encounters between rare mutants. The relation between inbreeding and the evolution of adaptive drive also stands in stark contrast to the suppressive effect of inbreeding on the spread of selfish drivers (Burt and Trivers 1999, Bull 2017).

The interaction of selfing and all-and-none segregation in the present work is consonant with Charlesworth’s (1979) model investigating the spread of sex-limited gamete lethals (or detrimentals) in a system of selfing and balanced homozygotic lethality, characteristics which are common in species with permanent translocation heterozygosity. Charlesworth found that the invasion of a sex-limited gamete lethal (in sperm, say) is effectively neutral under full selfing, but disadvantageous at lower selfing rates; subsequent invasion of an opposite-sex gamete lethal (i.e. lethal eggs of the other allelic type) is selectively favored since this would avoid production of lethal homozygous progeny; the all-and-none scheme could thus become fixed. Holsinger and Ellstrand (1984) remark that an even less restrictive picture would emerge if the gamete lethals of Charlesworth’s model were replaced with meiotic drivers.

Following Eshel (1985) and Úbeda and Haig (2005), here I adopt the modifier’s eye view of long-term evolution, rather than focusing on the short-term dynamics of meiotic drive alleles per se. The modifier perspective is key, since whatever the short-term evolutionary dynamics of any particular driver may be, the ultimate fate of drivers over the long-term will owe to the make-up of the genetic background (Eshel 1996, Eshel and Feldman 2001). If the genes capable of modifying drive phenotypes overwhelmingly benefit from suppressing such behavior, then any particular meiotic driver is destined for extinction on a timescale that is limited only by the introduction and spread of suppressor alleles at the relevant loci. Given that unlinked modifiers are likely to be more numerous than linked modifiers from the perspective of any particular distorter, the dissent or approval of the permanent unlinked majority of the parliament of genes is decisive for determining the long run stability of the drive phenotype. With non-zero rates of self-fertilization, the unlinked majority ought to favor the repeal of Mendelian law at any locus that is at a stable equilibrium due to classical heterozygote advantage. Under random mating, this expectation holds for asymmetric overdominance, but only mild assent is expected for the classical symmetric case (Úbeda and Haig 2005). It remains to be discovered whether there exist other classes of initial population conditions that effect the invasion of unlinked drive-enhancers.

Given the ubiquity of Mendelian segregation in nature, how come real populations do not conform to theoretical expectation? One possibility is suggested in the numerical results, which indicate that a concatenation of weak coefficients in the selection and segregation processes fails to effect the swift establishment of a mutant modifier. The lifespan of a balanced fitness polymorphism that is the target of adaptive drive presumably sets a time limit during which the appropriate modifier mutations must originate, survive genetic drift at the extinction boundary, and then rise from rarity to an appreciable frequency. If typical balanced polymorphisms do not last long enough for the full course of evolutionary changes needed for the establishment of such modifiers, then the ubiquity of Mendelian segregation follows. In allowing for the occasional case of long-persisting overdominance, such a constraint permits the odd example of adaptive drive as in the permanent translocation heterozygotes of *Oenothera* that adhere to the all-and-none scheme in gametogenesis (Cleland 1972, Holsinger and Ellstrand 1984, Harte 1994). A related issue is that heterozygote advantage might often be associated with rapid allelic turnover (Sellis et al. 2011) in a manner that interferes with an orderly process of modifier selection. Furthermore, as some of the numerical examples of asymmetrically-reflected drive ratios reveal, some mutants can knock out the polymorphism of heterotic alleles, and in doing so collapse the basis for their own stable persistence.

As mentioned by Úbeda and Haig (2005), male fertility costs associated with segregation distorters can also be a serious constraint (Haig and Bergstrom 1995). For selfers, the costs of reduced male fertility under adaptive drive would strongly depend on the intensity of inter-individual male gamete competition. Such competition is presumably less intense for predominant selfers, assuming a negative relationship between the rate of selfing and population density (Karron et al. 1995, Morgan, Wilson and Knight 2005). Populations with intermediate and low selfing rates, however, would indeed seem to be severely affected by this constraint.

Additionally, there is a lack of realism associated with having a single modifier directly control the production of both male and female gametes (Úbeda 2006). An attempt to adhere closely to realism would involve investigating a rather complicated model that incorporates two sex-limited modifier genes encoded at different loci (one for each sex), as well as multiple alleles at the fitness locus representing the simultaneous variation at *A* of sex-limited distorters and wild-type Mendelian alleles. It is not clear whether the simplifying assumption on modifier control is ultimately misleading.

While these considerations and others may be relevant, at present a decisive argument for why all-and-none segregation is not the normal mode of inheritance in natural populations, selfing or otherwise, has not been established. The near-universality of Mendelian segregation rests securely on observation but lacks a convincing adaptationist explanation. Perhaps a conclusive idea of the pertinent constraints will emerge from more realistic multi-locus scenarios. What is clear is that our understanding has shifted from the feeling of a satisfying denouement derived from earlier sex-independent models to a sort of puzzlement that stems from consideration of sex-differentiated meiosis.

## Data accessibility

A Mathematica notebook (Wolfram Research, Inc. 2018, ver 11.2) and an R script (R Core Team 2018) for numerical iteration are available at: https://github.com/ebrud/adaptive_meiotic_drive.

## Acknowledgements

I thank Walter Eanes, Mark Kirkpatrick, John True, and Joshua Rest for helpful comments on the manuscript.

## Appendix

Explanation of the **T** matrix: the parent genotype indicated on the row label produces the offspring indicated on the column labels, according to the proportions given by the elements. The elements incorporate (1) selection, (2) free recombination, and (3) arbitrary segregation parameters. The matrix is comparable to that of Karlin (1968, Table 4, p. 246), who assumed equal segregation and arbitrary recombination.

(The **T** matrix describes the production of offspring due to selfing. If random mating also occurs (i.e. mixed mating, 0 < S < 1), then the **U** and **V** matrices describing outcrossing are also involved. See the description of the Model in the main text).

Explanation of the **U** and **V** matrices: the genotypes on the row labels produce the gametes indicated on the column labels in the proportions given by the elements, incorporating processes 1-3 as in the paragraph above. See next page.

While the **T** matrix directly figures in an analytical derivation (see the section on obligate selfing), the **U** and **V** matrices are merely convenient references for the compact representation of the system of equations given in the Model section.

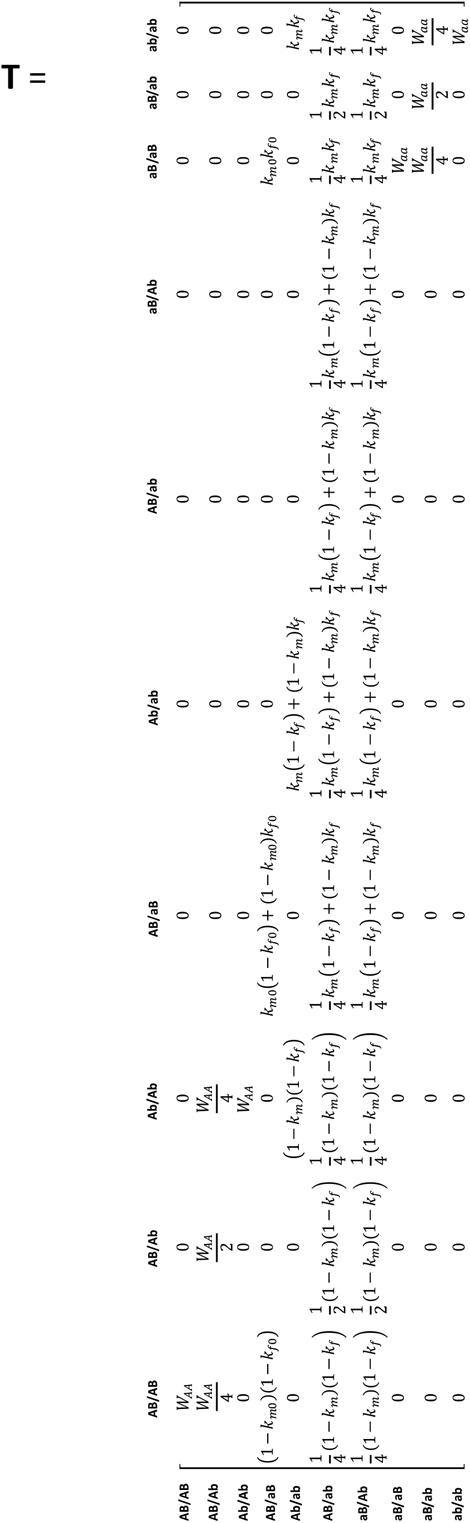

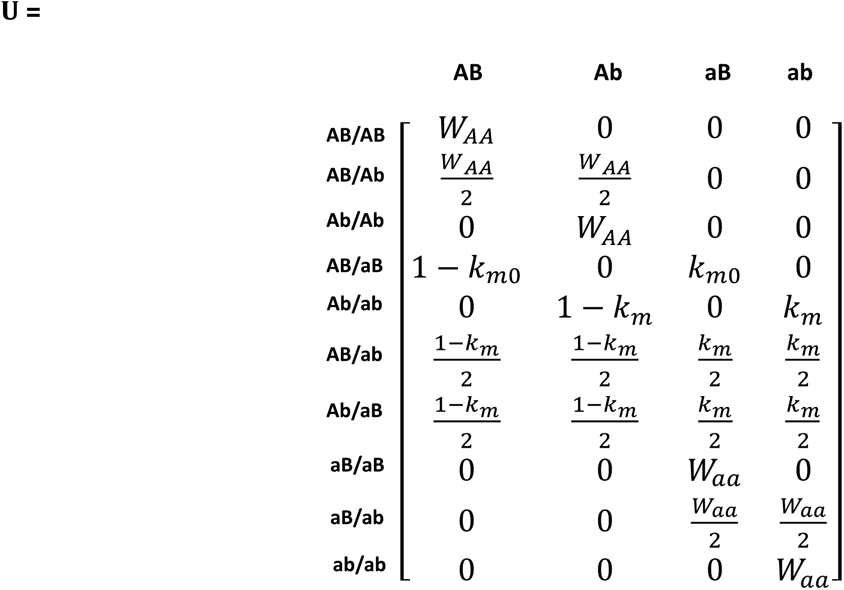

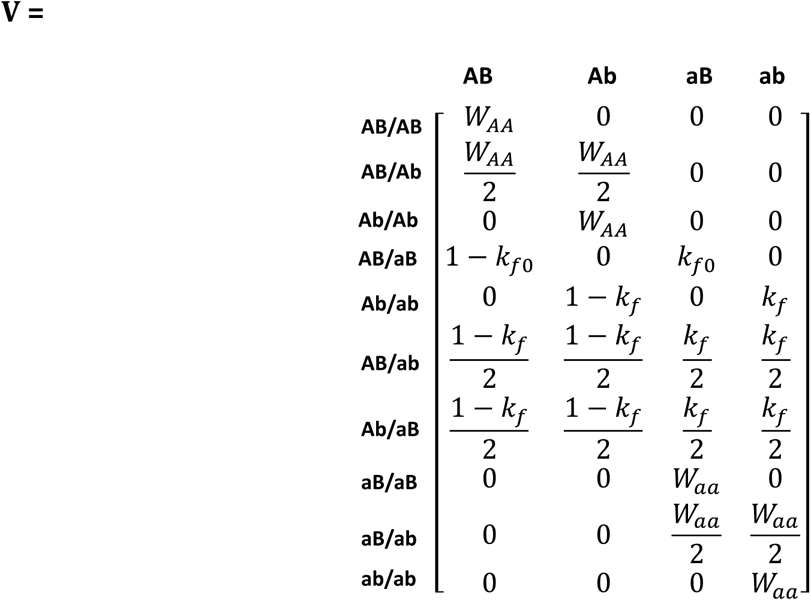

