## Supplemental file for "Adaptive meiotic drive in selfing populations with heterozygote advantage"

Evgeny Brud

### S1) Jacobian matrix:

For **Result 1** and **Result 2**, the Jacobian matrix is given by:

$$\begin{matrix}
\frac{\partial p'_{11}}{\partial p_{11}} & \frac{\partial p'_{11}}{\partial p_{33}} & \frac{\partial p'_{11}}{\partial p_{13}} & \frac{\partial p'_{11}}{\partial p_{12}} & \frac{\partial p'_{11}}{\partial p_{14}} & \frac{\partial p'_{11}}{\partial p_{23}} & \frac{\partial p'_{11}}{\partial p_{34}} & \frac{\partial p'_{11}}{\partial p_{22}} & \frac{\partial p'_{11}}{\partial p_{24}} & \frac{\partial p'_{11}}{\partial p_{44}} \\
\frac{\partial p'_{33}}{\partial p_{11}} & \frac{\partial p'_{33}}{\partial p_{33}} & \frac{\partial p'_{33}}{\partial p_{13}} & \frac{\partial p'_{33}}{\partial p_{12}} & \frac{\partial p'_{33}}{\partial p_{14}} & \frac{\partial p'_{33}}{\partial p_{23}} & \frac{\partial p'_{33}}{\partial p_{34}} & \frac{\partial p'_{33}}{\partial p_{22}} & \frac{\partial p'_{33}}{\partial p_{24}} & \frac{\partial p'_{33}}{\partial p_{44}} \\
\frac{\partial p'_{13}}{\partial p_{11}} & \frac{\partial p'_{13}}{\partial p_{33}} & \frac{\partial p'_{13}}{\partial p_{13}} & \frac{\partial p'_{13}}{\partial p_{12}} & \frac{\partial p'_{13}}{\partial p_{14}} & \frac{\partial p'_{13}}{\partial p_{23}} & \frac{\partial p'_{13}}{\partial p_{34}} & \frac{\partial p'_{13}}{\partial p_{22}} & \frac{\partial p'_{13}}{\partial p_{24}} & \frac{\partial p'_{13}}{\partial p_{44}} \\
\frac{\partial p'_{12}}{\partial p_{11}} & \frac{\partial p'_{12}}{\partial p_{33}} & \frac{\partial p'_{12}}{\partial p_{13}} & \frac{\partial p'_{12}}{\partial p_{12}} & \frac{\partial p'_{12}}{\partial p_{14}} & \frac{\partial p'_{12}}{\partial p_{23}} & \frac{\partial p'_{12}}{\partial p_{34}} & \frac{\partial p'_{12}}{\partial p_{22}} & \frac{\partial p'_{12}}{\partial p_{24}} & \frac{\partial p'_{12}}{\partial p_{44}} \\
\frac{\partial p'_{14}}{\partial p_{11}} & \frac{\partial p'_{14}}{\partial p_{33}} & \frac{\partial p'_{14}}{\partial p_{13}} & \frac{\partial p'_{14}}{\partial p_{12}} & \frac{\partial p'_{14}}{\partial p_{14}} & \frac{\partial p'_{14}}{\partial p_{23}} & \frac{\partial p'_{14}}{\partial p_{34}} & \frac{\partial p'_{14}}{\partial p_{22}} & \frac{\partial p'_{14}}{\partial p_{24}} & \frac{\partial p'_{14}}{\partial p_{44}} \\
\frac{\partial p'_{23}}{\partial p_{11}} & \frac{\partial p'_{23}}{\partial p_{33}} & \frac{\partial p'_{23}}{\partial p_{13}} & \frac{\partial p'_{23}}{\partial p_{12}} & \frac{\partial p'_{23}}{\partial p_{14}} & \frac{\partial p'_{23}}{\partial p_{23}} & \frac{\partial p'_{23}}{\partial p_{34}} & \frac{\partial p'_{23}}{\partial p_{22}} & \frac{\partial p'_{23}}{\partial p_{24}} & \frac{\partial p'_{23}}{\partial p_{44}} \\
\frac{\partial p'_{34}}{\partial p_{11}} & \frac{\partial p'_{34}}{\partial p_{33}} & \frac{\partial p'_{34}}{\partial p_{13}} & \frac{\partial p'_{34}}{\partial p_{12}} & \frac{\partial p'_{34}}{\partial p_{14}} & \frac{\partial p'_{34}}{\partial p_{23}} & \frac{\partial p'_{34}}{\partial p_{34}} & \frac{\partial p'_{34}}{\partial p_{22}} & \frac{\partial p'_{34}}{\partial p_{24}} & \frac{\partial p'_{34}}{\partial p_{44}} \\
\frac{\partial p'_{22}}{\partial p_{11}} & \frac{\partial p'_{22}}{\partial p_{33}} & \frac{\partial p'_{22}}{\partial p_{13}} & \frac{\partial p'_{22}}{\partial p_{12}} & \frac{\partial p'_{22}}{\partial p_{14}} & \frac{\partial p'_{22}}{\partial p_{23}} & \frac{\partial p'_{22}}{\partial p_{34}} & \frac{\partial p'_{22}}{\partial p_{22}} & \frac{\partial p'_{22}}{\partial p_{24}} & \frac{\partial p'_{22}}{\partial p_{44}} \\
\frac{\partial p'_{24}}{\partial p_{11}} & \frac{\partial p'_{24}}{\partial p_{33}} & \frac{\partial p'_{24}}{\partial p_{13}} & \frac{\partial p'_{24}}{\partial p_{12}} & \frac{\partial p'_{24}}{\partial p_{14}} & \frac{\partial p'_{24}}{\partial p_{23}} & \frac{\partial p'_{24}}{\partial p_{34}} & \frac{\partial p'_{24}}{\partial p_{22}} & \frac{\partial p'_{24}}{\partial p_{24}} & \frac{\partial p'_{24}}{\partial p_{44}} \\
\frac{\partial p'_{44}}{\partial p_{11}} & \frac{\partial p'_{44}}{\partial p_{33}} & \frac{\partial p'_{44}}{\partial p_{13}} & \frac{\partial p'_{44}}{\partial p_{12}} & \frac{\partial p'_{44}}{\partial p_{14}} & \frac{\partial p'_{44}}{\partial p_{23}} & \frac{\partial p'_{44}}{\partial p_{34}} & \frac{\partial p'_{44}}{\partial p_{22}} & \frac{\partial p'_{44}}{\partial p_{24}} & \frac{\partial p'_{44}}{\partial p_{44}}
\end{matrix}$$

The Jacobian matrix above is evaluated at a resident Mendelian equilibrium (**Result 1**) or a resident all-and-none equilibrium (**Result 2**). Evaluation at either state yields a block matrix of the form:

$$\mathbf{J}|_{(\hat{q},\hat{F})} = \begin{matrix} \mathbf{J_{res}} & \mathbf{V} \\ \mathbf{0} & \mathbf{J_{mut}} \end{matrix}$$

The lower left submatrix is a 7x3 block of zeroes, and so the stability of the system depends solely on the eigenvalues of the square block matrices on the diagonal, $\mathbf{J_{res}}$ and $\mathbf{J_{mut}}$. (See Otto and Day 2007 for the general theory on evolutionary invasion analysis).

For **Result 1** (external stability of a Mendelian population), the corresponding submatrices, $\mathbf{J_{res}}$ and $\mathbf{J_{mut}}$, are given by:

y$\mathbf{J}_{\mathbf{res}}$ =

$$\begin{pmatrix} W_G\left(-W_G S^2+\left(2W_G^2+W_G-Z-1\right)S+2(W_G-1)(Z+W_G+1)\right) & -W_G\left(W_G S^2+\left(2W_G^2-5W_G+Z+1\right)S+2(W_G-1)(Z+W_G+1)\right) & -S W_G(Z+S W_G+W_G-3) \\ -W_G\left(W_G S^2+\left(2W_G^2-5W_G+Z+1\right)S+2(W_G-1)(Z+W_G+1)\right) & W_G\left(-W_G S^2+\left(2W_G^2+W_G-Z-1\right)S+2(W_G-1)(Z+W_G+1)\right) & -S W_G(Z+S W_G+W_G-3) \\ 2S W_G(Z+(S-3)W_G+1) & 2S W_G(Z+(S-3)W_G+1) & 2S W_G(Z+S W_G+W_G-3) \end{pmatrix}$$

x$\mathbf{J}_{\mathbf{mut}}$ =

$$\begin{matrix} 2W_G & -(S+1)k_m+k_f(S(2k_m-1)-1)+2 & -(S+1)k_m+k_f(S(2k_m-1)-1)+2 & 0 & -4(S-1)W_G & 2(S-1)(k_f+k_m-2) & 0 \\ 0 & -2S k_m k_f+k_f+k_m & -2S k_m k_f+k_f+k_m & -2(S-1)W_G & 0 & -2(S-1)(k_f+k_m) & -4(S-1)W_G \\ -2(S-1)W_G & -2S+k_f(-2S(k_m-1)-1)+(2S-1)k_m+2 & -2S+k_f(-2S(k_m-1)-1)+(2S-1)k_m+2 & 0 & -4(S-1)W_G & 2(S-1)(k_f+k_m-2) & 0 \\ 0 & S(2k_m-1)k_f+k_f-S k_m+k_m & S(2k_m-1)k_f+k_f-S k_m+k_m & 2W_G & 0 & -2(S-1)(k_f+k_m) & -4(S-1)W_G \\ S W_G & S(k_f-1)(k_m-1) & S(k_f-1)(k_m-1) & 0 & 4S W_G & 4S(k_f-1)(k_m-1) & 0 \\ 0 & S(-2k_m k_f+k_f+k_m) & S(-2k_m k_f+k_f+k_m) & 0 & 0 & 4S(-2k_m k_f+k_f+k_m) & 0 \\ 0 & S k_f k_m & S k_f k_m & S W_G & 0 & 4S k_f k_m & 4S W_G \end{matrix}$$

, where x = $S W_G + W_G + Z + 1$

, y = $(W_G - 1)(S W_G + W_G + Z + 1)^2$

and Z = $\sqrt{(S+1)^2 W_G^2 + (2 - 6S) W_G + 1}$

---

For **Result 2** (external stability of an all-and-none population), the corresponding submatrices, $\mathbf{J}_{\mathbf{res}}$ and $\mathbf{J}_{\mathbf{mut}}$, are given by:

$\mathbf{J}_{\mathbf{res}}$ = $$\begin{pmatrix} W_G & 0 & 0 \\ 0 & W_G & 0 \\ -W_G & -W_G & 0 \end{pmatrix}$$

and

$\mathbf{J}_{\mathbf{mut}}$ =

$$\begin{pmatrix} \frac{1}{2}(S-3)W_G & \frac{1}{4}(3k_m S+2S-2k_m-k_f(2k_m S+S-2)-6) & \frac{1}{4}(3k_m S+2S-2k_m-k_f(2k_m S+S-2)-6) & \frac{1}{2}(S-3)W_G & (S-2)W_G & S-2 & (S-2)W_G \\ \frac{W_G}{2} & \frac{1}{2}(k_f-1)(S k_m-1) & \frac{1}{2}(k_f-1)(S k_m-1) & 0 & W_G-S W_G & (S-1)(k_f-1) & 0 \\ 0 & \frac{1}{4}(S k_m-k_f(2k_m S+S-2)) & \frac{1}{4}(S k_m-k_f(2k_m S+S-2)) & -\frac{1}{2}(S-1)W_G & 0 & k_f-S k_f & W_G-S W_G \\ -\frac{1}{2}(S-1)W_G & \frac{1}{4}(S(k_f-2)+(S(3-2k_f)-2)k_m+2) & \frac{1}{4}(S(k_f-2)+(S(3-2k_f)-2)k_m+2) & 0 & W_G-S W_G & (S-1)(k_m-1) & 0 \\ 0 & \frac{1}{2}(S(k_f-1)k_m+k_m) & \frac{1}{2}(S(k_f-1)k_m+k_m) & \frac{W_G}{2} & 0 & k_m-S k_m & W_G-S W_G \\ \frac{S W_G}{4} & \frac{1}{4}S(k_f-1)(k_m-1) & \frac{1}{4}S(k_f-1)(k_m-1) & 0 & S W_G & S(k_f-1)(k_m-1) & 0 \\ 0 & -\frac{1}{4}S(k_f(2k_m-1)-k_m) & -\frac{1}{4}S(k_f(2k_m-1)-k_m) & 0 & 0 & S(-2k_m k_f+k_f+k_m) & 0 \\ 0 & \frac{1}{4}S k_f k_m & \frac{1}{4}S k_f k_m & \frac{S W_G}{4} & 0 & S k_f k_m & S W_G \end{pmatrix}$$

### S2) Stability analysis of a single-locus internal Mendelian equilibrium under partial selfing and heterozygote advantage:

$(\hat{q}, \hat{F})$ under Mendelian segregation is provided in the text and is internally stable under partial selfing and heterozygote advantage if the eigenvalues of the following Jacobian matrix (evaluated at equilibrium) are all less than one in absolute value:

$$\left.\begin{matrix} \frac{\partial q'}{\partial q} & \frac{\partial q'}{\partial F} \\ \frac{\partial F'}{\partial q} & \frac{\partial F'}{\partial F} \end{matrix}\right|_{(\hat{q},\hat{F})} = \begin{pmatrix} \frac{4\mathrm{W}_G}{(S+1)\mathrm{W}_G+\sqrt{(S+1)^2\mathrm{W}_G{}^2+(2-6S)\mathrm{W}_G+1}+1} & 0 \\ 0 & \frac{8S\mathrm{W}_G}{\left((S+1)\mathrm{W}_G+\sqrt{(S+1)^2\mathrm{W}_G{}^2+(2-6S)\mathrm{W}_G+1}+1\right)^2} \end{pmatrix}.$$

The eigenvalues are the elements on the diagonal. Stability conditions correspond to any amount of partial selfing and any amount of heterozygote advantage (here, assuming symmetric selection against homozygotes).

---

### S3) Stability analysis of single-locus internal all-and-none equilibrium under partial selfing and heterozygote advantage:

Under all-and-none segregation, $(\hat{q}, \hat{F})$ equals (1/2, -1) and is internally stable for partial selfing and heterozygote advantage if the eigenvalues of the following Jacobian matrix are all less than one in absolute value:

$$\left.\begin{matrix} \frac{\partial q'}{\partial q} & \frac{\partial q'}{\partial F} \\ \frac{\partial F'}{\partial q} & \frac{\partial F'}{\partial F} \end{matrix}\right|_{(\hat{q},\hat{F})} = \begin{pmatrix} \mathrm{W}_G & 0 \\ 0 & \mathrm{W}_G \end{pmatrix}$$

The eigenvalues are both $W_G$. Stability conditions correspond to any amount of symmetric selection against homozygotes.

With generalized homozygote fitnesses,

$$\left.\begin{matrix} \frac{\partial q'}{\partial q} & \frac{\partial q'}{\partial F} \\ \frac{\partial F'}{\partial q} & \frac{\partial F'}{\partial F} \end{matrix}\right|_{(\hat{q},\hat{F})} = \begin{pmatrix} \frac{\mathrm{W_{aa}} + \mathrm{W_{AA}}}{2} & \frac{\mathrm{W_{aa}} - \mathrm{W_{AA}}}{8} \\ 2(\mathrm{W_{aa}} - \mathrm{W_{AA}}) & \frac{\mathrm{W_{aa}} + \mathrm{W_{AA}}}{2} \end{pmatrix}$$

The eigenvalues are $W_{AA}$ and $W_{aa}$, with stability requiring each homozygote fitness to be less than 1 (i.e. any amount of heterozygote advantage).

---

### S4) Stability analysis of single-locus internal Mendelian equilibrium under full selfing and generalized heterozygote advantage:

Assuming full selfing (S=1) and generalized homozygote fitnesses, stability requires a greater than two-fold heterozygote advantage over each homozygote. Solving for equilibrium:

$$\hat{q} = \frac{(\mathrm{W_{aa}} - 1)(2\mathrm{W_{AA}} - 1)}{\mathrm{W_{aa}}(4\mathrm{W_{AA}} - 3) - 3\mathrm{W_{AA}} + 2}$$

$$\hat{F} = \frac{\mathrm{W_{AA}} + \mathrm{W_{aa}} - 2\mathrm{W_{AA}}\mathrm{W_{aa}}}{2\mathrm{W_{AA}}\mathrm{W_{aa}} - 2\mathrm{W_{AA}} - 2\mathrm{W_{aa}} + 2}$$

$$\left.\begin{matrix} \frac{\partial q'}{\partial q} & \frac{\partial q'}{\partial F} \\ \frac{\partial F'}{\partial q} & \frac{\partial F'}{\partial F} \end{matrix}\right|_{(\hat{q},\hat{F})} = \begin{pmatrix} \frac{2\left((2 W_{AA}-1) W_{aa}^2+\left(2 W_{AA}^2-4 W_{AA}+1\right) W_{aa}-(W_{AA}-1) W_{AA}\right)}{-3 W_{AA}+W_{aa} (4 W_{AA}-3)+2} & \frac{4 (W_{aa}-1)^2 (2 W_{aa}-1) (W_{aa}-W_{AA}) (W_{AA}-1)^2 (2 W_{AA}-1)}{(-3 W_{AA}+W_{aa} (4 W_{AA}-3)+2)^3} \\ \frac{(W_{aa}-W_{AA}) (-3 W_{AA}+W_{aa} (4 W_{AA}-3)+2)}{(W_{aa}-1) (W_{AA}-1)} & \frac{2\left(2 (W_{AA}-1) W_{aa}^2+\left(2 W_{AA}^2-2 W_{AA}+1\right) W_{aa}-2 W_{AA}^2+W_{AA}\right)}{-3 W_{AA}+W_{aa} (4 W_{AA}-3)+2} \end{pmatrix}$$

The eigenvalues are $2\mathrm{W_{AA}}$ and $2\mathrm{W_{aa}}$, which are obviously less than one only if both homozygote fitnesses are less than one-half. Hence, full selfing requires a greater than two-fold heterozygote advantage in order to establish a stable internal equilibrium at A.

---

### S5) Invasion analysis of complete drive-suppressors in an all-and-none resident population with generalized homozygote fitnesses

Given that a resident population is fixed for all-and-none segregation, it is possible to ascertain whether the direct invasion of a drive suppressor will occur for generalized homozygous fitnesses. There are two scenarios of interest:

1) Direct invasion of a sex-limited drive-suppressor that imposes Mendelian segregation but maintains the resident maximal distortion in the opposite sex. (e.g. a female-limited suppressor that restores equal segregation in eggs, but doesn't affect male meiosis).

2) Direct invasion of a sex-independent drive-suppressor which imposes Mendelian segregation in both sexes, thereby abolishing sex-specific meiotic drive altogether. (Perhaps less realistic than the first scenario, but it is also tractable).

For scenario 1 (sex-limited restoration of Mendelian segregation), the eigenvalues are:

$$\lambda_{S5,1,1} = 0$$

$$\lambda_{S5,1,2} = \frac{S}{4}$$

$$\lambda_{S5,1,3} = \frac{S\mathrm{W}_{\mathrm{aa}}}{2}$$

$$\lambda_{S5,1,4} = \frac{1}{2}(1+S)\mathrm{W}_{\mathrm{aa}}$$

$$\lambda_{S5,1,5} = \frac{S\mathrm{W}_{\mathrm{AA}}}{2}$$

$$\lambda_{S5,1,6} = \frac{1}{8}\left(3 - S + 2\mathrm{W}_{\mathrm{AA}} + 2S\mathrm{W}_{\mathrm{AA}} - \frac{1}{2}\sqrt{-64(1+S)\mathrm{W}_{\mathrm{AA}} + 4(3 + 2\mathrm{W}_{\mathrm{AA}} + S(2\mathrm{W}_{\mathrm{AA}} - 1)^2)}\right)$$

$$\lambda_{S5,1,7} = \frac{1}{8}\left(3 - S + 2\mathrm{W}_{\mathrm{AA}} + 2S\mathrm{W}_{\mathrm{AA}} + \frac{1}{2}\sqrt{-64(1+S)\mathrm{W}_{\mathrm{AA}} + 4(3 + 2\mathrm{W}_{\mathrm{AA}} + S(2\mathrm{W}_{\mathrm{AA}} - 1)^2)}\right)$$

These eigenvalues are uniformly less than one in magnitude for any heterozygote advantage and partial selfing. All-and-none is therefore stable to such “large-effect” sex-limited suppressors.

For scenario 2 (sex-independent restoration of Mendelian segregation), the eigenvalues are complicated, and are available in compact form only as “Root” objects in Mathematica. The reader is encouraged to consult the Mathematica notebook attached to this article to see that the eigenvalues lie between -1 and 1 for scenario 2.

---

### S6) Deriving the optimal rate of selfing that maximizes the rate of modifier invasion into a resident Mendelian population:

Because the rate of invasion of a rare modifier into a Mendelian population is given by leading eigenvalue of $\mathbf{J}_{\mathbf{mut}}$ ($\lambda_L$, Eq. 3, main text), the partial derivative of $\lambda_L$ with respect to S (the selfing parameter) can be set equal to zero, and subsequently solved for an expression of S that maximizes the value of $\lambda_L$ (which, as demonstrated in the Mathematica file, never attains negative or complex values). This expression for S is termed $S_{opt}$ (as in ‘optimal’) below.

The equation to solve for $S_{opt}$ is given by:

$$\frac{\partial \lambda_L}{\partial S} = 0$$

This yields a conditional expression for $S_{opt}$ equal to

$$\frac{2(W_G+1)}{3W_G+4k^2-4k+1}-\sqrt{\frac{-4k^2W_G^2-8k^2W_G+4kW_G^2+8kW_G+W_G^3+W_G^2-W_G-4k^2+4k-1}{W_G\left(3W_G+4k^2-4k+1\right)^2}}$$

, which is valid as long as segregation is non-Mendelian and $\frac{-1}{4k^2-4k-1} < W_G < 1$ (Fig. S2).

### Supplementary Figures

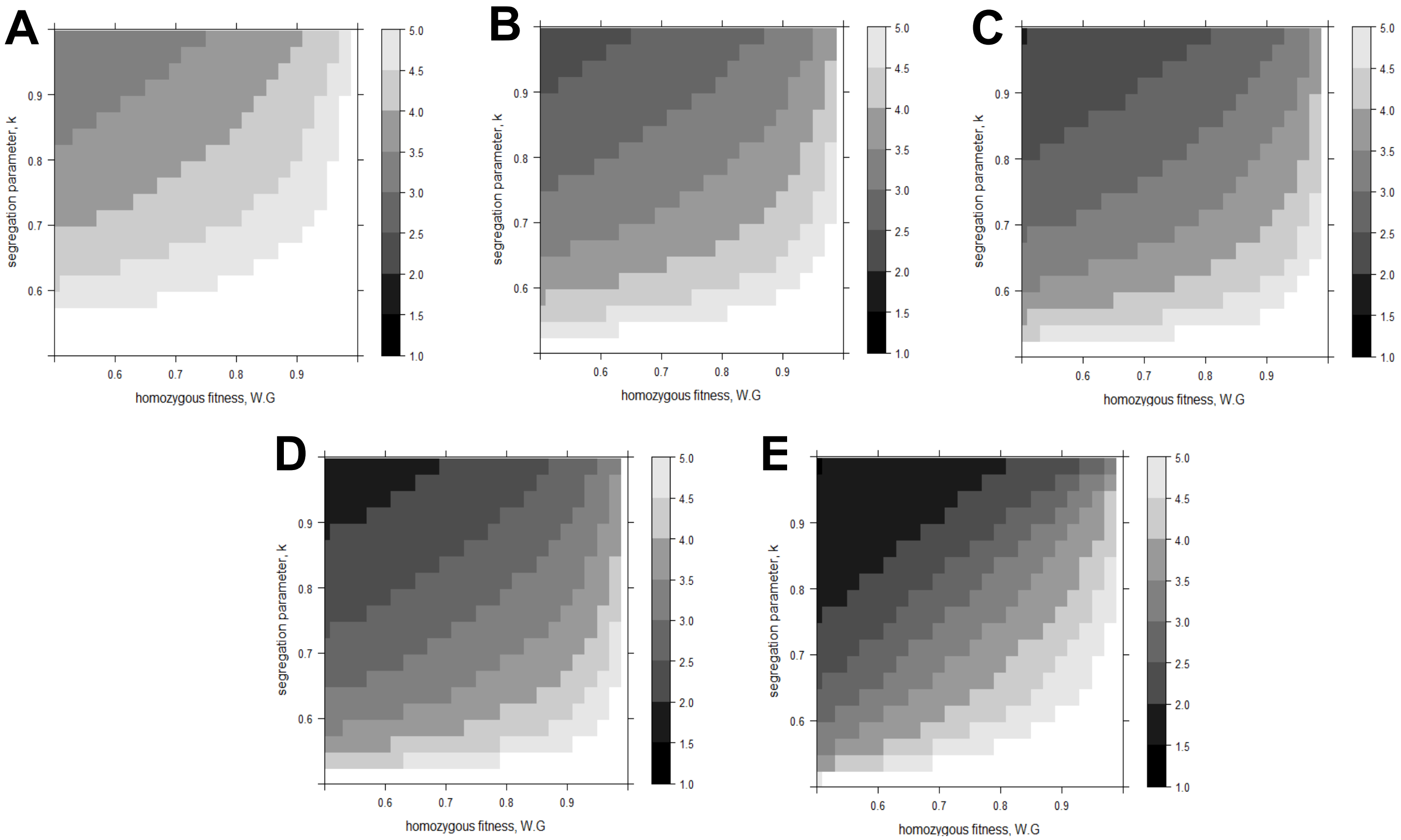


Fig. S1. **Time required for adaptive drive enhancer "fixation" in a resident Mendelian population for various parameter combinations.** (A-E) Each panel is a heatmap that depicts the time (in $\log_{10}$ generations) that it takes for a modifier (b allele) to reach 99% frequency, after being introduced at an initial frequency of 0.01% into a resident Mendelian population at an internal heterotic equilibrium at A (assuming initial linkage equilibrium). The heatmaps depict the outcome of numerical iterations run for $10^5$ generations in total for perfectly sex-reflected segregation schemes and symmetric selection against homozygotes. Darker areas correspond to faster rates of b allele "fixation", while white areas correspond to parameter combinations in which the b allele failed to "fix" within $10^5$ generations. Each heatmap consists of 500 parameter combinations, composed of 25 homozygous fitness values (from $W_G = 0.5$ to $W_G = 0.98$ separated by increments of 0.02) and 20 segregation parameter values (from k = 0.51 to k = 0.985 separated by increments of 0.025). The selfing rates (S) are: (A) 2.5%, (B) 26.25%, (C) 50%, (D) 73.75%, (E) 97.5%. The non-monotonicity of the relation between selfing rates and the rate of modifier fixation can be noticed, for instance, by comparing panels D and E (e.g. comparing the area of the white region).

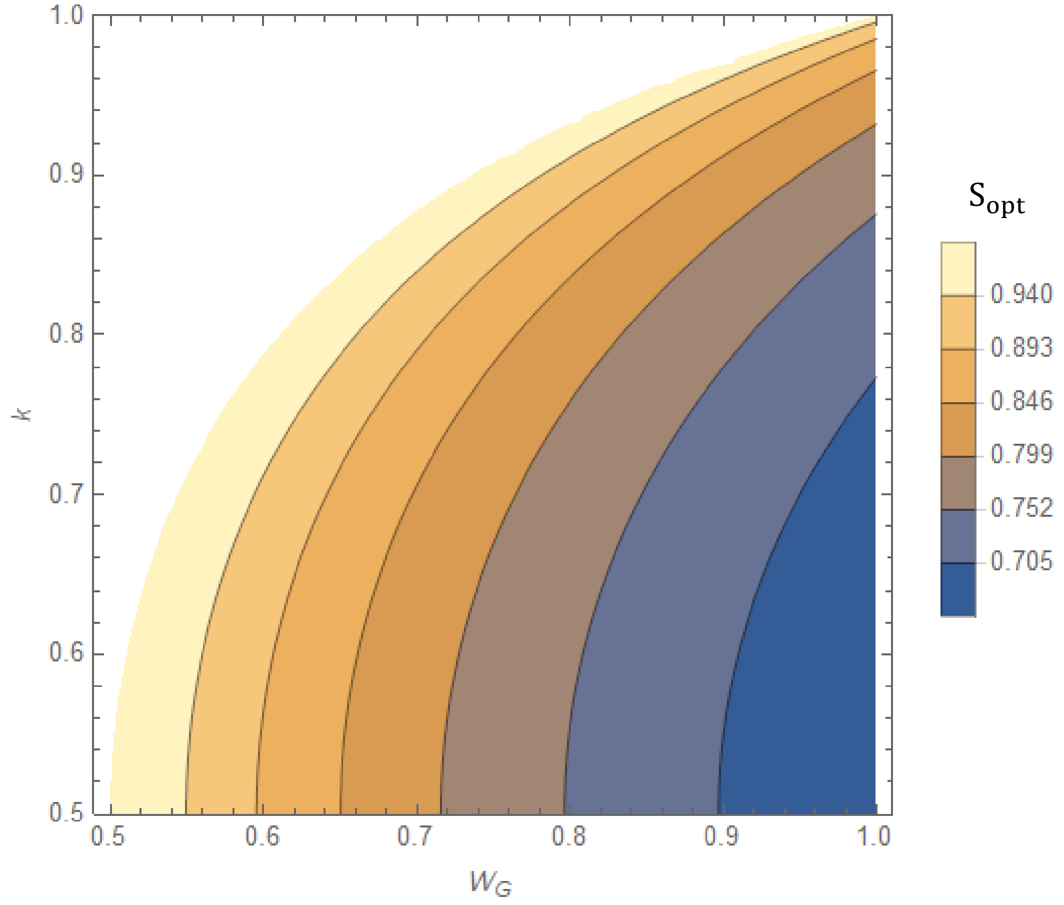


Fig. S2. **Optimal selfing rates ($S_{opt}$) for maximizing the rate of modifier invasion into a resident Mendelian population.** The contour plot depicts the value of the selfing parameter that maximizes the leading eigenvalue (Eq. 3, main text) which is equal to the rate of adaptive drive enhancer (b allele) invasion into a resident Mendelian population. See section 5 of the Supplemental file for more information.
